# Compartment-driven imprinting of intestinal CD4 (regulatory) T cells in inflammatory bowel disease and homeostasis

**DOI:** 10.1101/2022.05.06.490870

**Authors:** Lisanne Lutter, José J.M. ter Linde, Eelco C. Brand, David P. Hoytema van Konijnenburg, Britt Roosenboom, Carmen Horjus Talabur-Horje, Bas Oldenburg, Femke van Wijk

## Abstract

**Objective:** The mucosal immune system is implicated in the etiology and progression of inflammatory bowel diseases. The lamina propria and epithelium of the gut mucosa constitute two separate compartments, containing distinct T cell populations. Human CD4 T cell programming and regulation of lamina propria and epithelium CD4 T cells, especially during inflammation, remains incompletely understood.

**Design:** We performed imaging mass cytometry, flow cytometry, bulk and single-cell RNA-sequencing to profile ileal lamina propria and intraepithelial CD4 T cells (CD4CD8αα, regulatory T cells (Tregs), CD69^-^ and CD69^high^ Trm T cells) in controls and Crohn’s disease (CD) patients (paired non-inflamed and inflamed).

**Results:** Inflammation results in alterations of the CD4 T cell population with a pronounced increase in Tregs and migrating/infiltrating cells. On a transcriptional level, inflammation within the epithelium induced T cell activation, increased IFNγ responses and effector Treg differentiation. Conversely, few transcriptional changes within the lamina propria were observed. Key regulators including the chromatin remodelers *ARID4B* and *SATB1* were found to drive compartment-specific transcriptional programming of CD4 T(reg) cells.

**Conclusion:** Inflammation in CD patients primarily induces changes within the epithelium and not the lamina propria. Additionally, there is compartment-specific CD4 T cell imprinting, driven by shared regulators, upon translocation from the lamina propria to the epithelium. The main consequence of epithelial translocation, irrespective of inflammation, seems to be an overall dampening of broad (pro-inflammatory) responses and tight regulation of lifespan. These data suggest differential regulation of the lamina propria and epithelium, with a specific regulatory role in the inflamed epithelium.

## Introduction

Inflammatory bowel disease (IBD), comprising Crohn’s disease (CD) and ulcerative colitis, is a chronic relapsing-remitting inflammatory disease with a multifactorial and not fully elucidated pathogenesis.^1^ Both the innate and adaptive immune system has been implicated in the etiology and progression of disease.^1^ The lamina propria and epithelium of the gut mucosa constitute two separate compartments, containing distinct populations of T cells.^2^ Intraepithelial T cells, interspersed between the single layer of epithelial cells, are in direct contact with food and microbiota-derived antigens, whereas T cells in the lamina propria are surrounded by a variety of other immune and stromal cells. In the lamina propria, most T cells are of the CD4 lineage, while CD8 T cells predominate in the epithelium.^3^ CD4 T cells are commonly divided in T helper (Th, effector) cells, regulatory T cells (Tregs) and CD4CD8αα T cells, with many of these being considered residents of the mucosal environment (tissue-resident memory T (Trm) cells, expressing CD69).^3,4^ In CD patients, CD4 T cells are increased in both the lamina propria and epithelium^5–8^ and changes of the phenotypic profile of several CD4 T cell subsets have been observed, including an increased percentage of (double/multiple) cytokine producing CD4 T cells (both regulatory and pro-inflammatory) and a more cytotoxic profile of these CD4 T cells.^9–14^ Additionally, it is now clear that microenvironmental cues can steer functional programs and cell dynamics that might have a role in health and disease. However, since studies reporting on CD4 T cells primarily focused on lamina propria or on the bulk of mucosal cells, our understanding of CD4 T cell programming in different mucosal compartments remains incompletely understood. This includes programming of Tregs and CD4CD8αα T cells, both associated with regulation,^15,16^ and the differential regulation of CD4 T cells derived from the lamina propria and epithelium, especially in inflammation. Studying tissue- and compartment-specific resident CD4 T cell regulation in IBD may improve our understanding of chronic and locally recurrent inflammation as well as open up new avenues for targeted treatment.

To assess these questions, we profiled surface marker-based CD4 T cell subsets with bulk RNA-sequencing as well as an unbiased CD4 T cell single-cell RNA-sequencing exploration, to map the CD4 T cell compartment in the epithelium and lamina propria of the human ileum, complemented with imaging mass cytometry (IMC), in control subjects and paired inflamed and non-inflamed ileal biopsies from patients with CD. Our results uncover compartment-specific CD4 T cell imprinting upon translocation from the lamina propria to the epithelium driven by key regulators regulating effector function and lifespan in the epithelium.

## Materials and Methods

### Participant inclusion

Patients with Crohn’s disease were prospectively enrolled at the Department of Gastroenterology and Hepatology, University Medical Center Utrecht and the outpatient clinic of the Rijnstate Crohn and Colitis Centre, Arnhem, The Netherlands. During ileocolonoscopy multiple biopsy specimens were taken for bulk (*n* = 4) and single-cell (*n* = 4) RNA-sequencing of sorted subsets and imaging mass cytometry (IMC; *n* = 3). Non-inflamed biopsies were taken > 2cm of macroscopically visible inflamed ileum. Control subjects for bulk RNA-sequencing (*n* = 3) and IMC (*n =* 2) did not have a diagnosis of IBD and underwent ileocolonoscopy for polyp surveillance and had normal macroscopical ileal mucosa (see table 1 for participant characteristics). Sex was not taken into account when selecting patients due to the sample size which would not allow for correction of the influence of sex on the immune system.

**Table 1.**
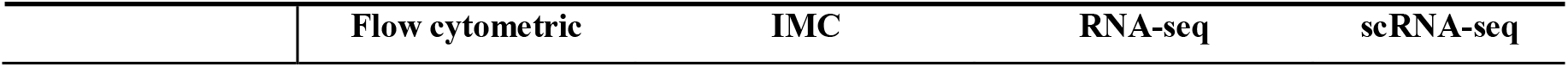
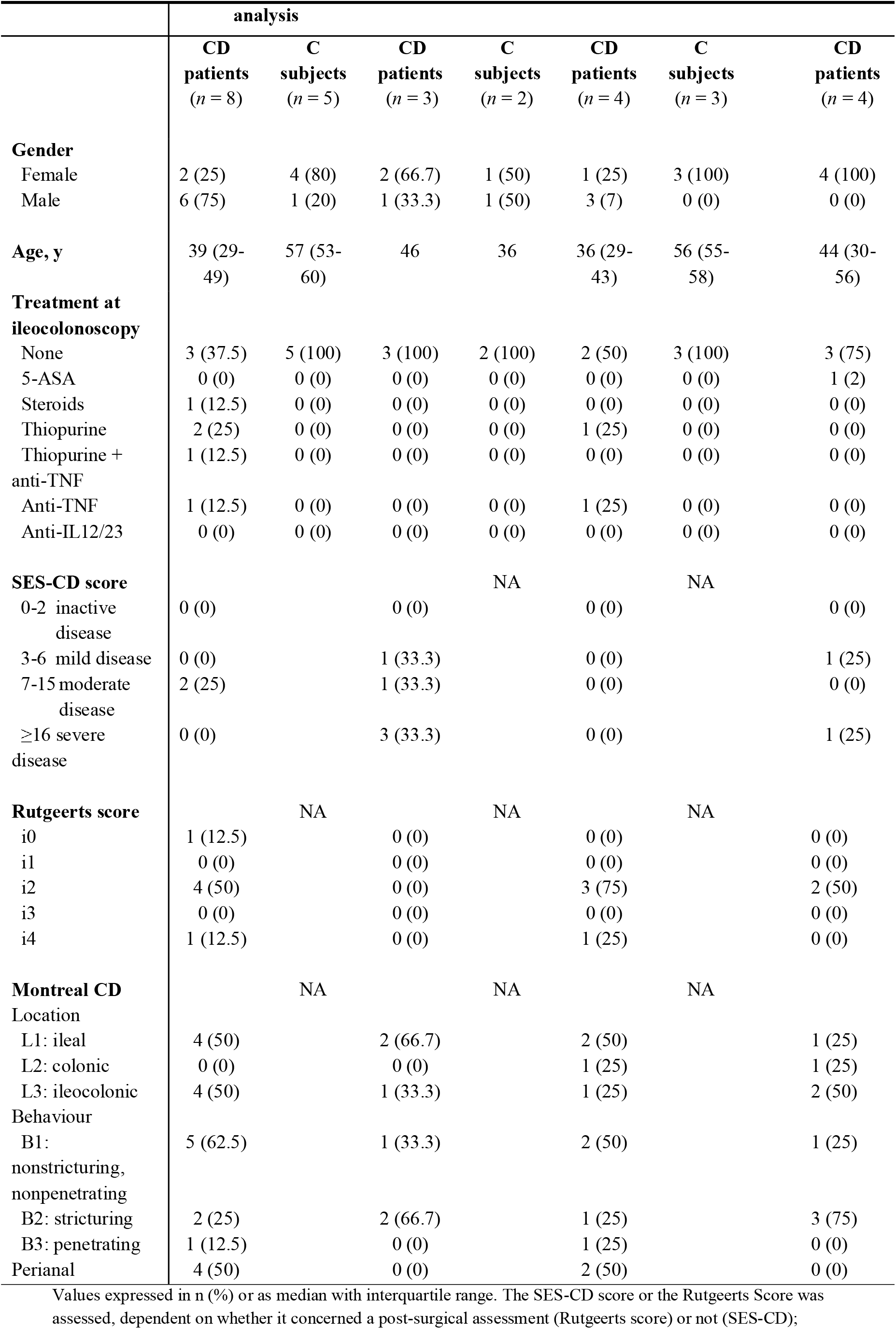
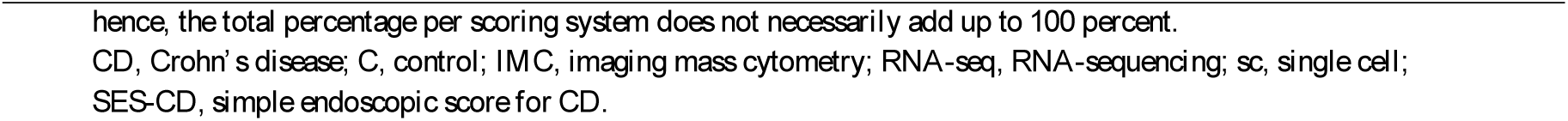
Baseline patient characteristics.

|  | Flow cytometric | IMC | RNA-seq | scRNA-seq |
| --- | --- | --- | --- | --- |

|  | analysis |  |  |  |  |  |  |
| --- | --- | --- | --- | --- | --- | --- | --- |
|  | CD<br>patients<br>(n = 8) | C<br>subjects<br>(n = 5) | CD<br>patients<br>(n = 3) | C<br>subjects<br>(n = 2) | CD<br>patients<br>(n = 4) | C<br>subjects<br>(n = 3) | CD<br>patients<br>(n = 4) |
| **Gender** |  |  |  |  |  |  |  |
| Female | 2 (25) | 4 (80) | 2 (66.7) | 1 (50) | 1 (25) | 3 (100) | 4 (100) |
| Male | 6 (75) | 1 (20) | 1 (33.3) | 1 (50) | 3 (7) | 0 (0) | 0 (0) |
| **Age, y** | 39 (29-49) | 57 (53-60) | 46 | 36 | 36 (29-43) | 56 (55-58) | 44 (30-56) |
| **Treatment at ileocolonoscopy** |  |  |  |  |  |  |  |
| None | 3 (37.5) | 5 (100) | 3 (100) | 2 (100) | 2 (50) | 3 (100) | 3 (75) |
| 5-ASA | 0 (0) | 0 (0) | 0 (0) | 0 (0) | 0 (0) | 0 (0) | 1 (2) |
| Steroids | 1 (12.5) | 0 (0) | 0 (0) | 0 (0) | 0 (0) | 0 (0) | 0 (0) |
| Thiopurine | 2 (25) | 0 (0) | 0 (0) | 0 (0) | 1 (25) | 0 (0) | 0 (0) |
| Thiopurine + anti-TNF | 1 (12.5) | 0 (0) | 0 (0) | 0 (0) | 0 (0) | 0 (0) | 0 (0) |
| Anti-TNF | 1 (12.5) | 0 (0) | 0 (0) | 0 (0) | 1 (25) | 0 (0) | 0 (0) |
| Anti-IL12/23 | 0 (0) | 0 (0) | 0 (0) | 0 (0) | 0 (0) | 0 (0) | 0 (0) |
| **SES-CD score** |  |  |  | NA |  | NA |  |
| 0-2 inactive disease | 0 (0) |  | 0 (0) |  | 0 (0) |  | 0 (0) |
| 3-6 mild disease | 0 (0) |  | 1 (33.3) |  | 0 (0) |  | 1 (25) |
| 7-15 moderate disease | 2 (25) |  | 1 (33.3) |  | 0 (0) |  | 0 (0) |
| ≥16 severe disease | 0 (0) |  | 3 (33.3) |  | 0 (0) |  | 1 (25) |
| **Rutgeerts score** |  | NA |  | NA |  | NA |  |
| i0 | 1 (12.5) |  | 0 (0) |  | 0 (0) |  | 0 (0) |
| i1 | 0 (0) |  | 0 (0) |  | 0 (0) |  | 0 (0) |
| i2 | 4 (50) |  | 0 (0) |  | 3 (75) |  | 2 (50) |
| i3 | 0 (0) |  | 0 (0) |  | 0 (0) |  | 0 (0) |
| i4 | 1 (12.5) |  | 0 (0) |  | 1 (25) |  | 0 (0) |
| **Montreal CD** |  | NA |  | NA |  | NA |  |
| **Location** |  |  |  |  |  |  |  |
| L1: ileal | 4 (50) |  | 2 (66.7) |  | 2 (50) |  | 1 (25) |
| L2: colonic | 0 (0) |  | 0 (0) |  | 1 (25) |  | 1 (25) |
| L3: ileocolonic | 4 (50) |  | 1 (33.3) |  | 1 (25) |  | 2 (50) |
| **Behaviour** |  |  |  |  |  |  |  |
| B1: nonstricturing, nonpenetrating | 5 (62.5) |  | 1 (33.3) |  | 2 (50) |  | 1 (25) |
| B2: stricturing | 2 (25) |  | 2 (66.7) |  | 1 (25) |  | 3 (75) |
| B3: penetrating | 1 (12.5) |  | 0 (0) |  | 1 (25) |  | 0 (0) |
| Perianal | 4 (50) |  | 0 (0) |  | 2 (50) |  | 0 (0) |

Values expressed in n (%) or as median with interquartile range. The SES-CD score or the Rutgeerts Score was assessed, dependent on whether it concerned a post-surgical assessment (Rutgeerts score) or not (SES-CD);
hence, the total percentage per scoring system does not necessarily add up to 100 percent. CD, Crohn's disease; C, control; IMC, imaging mass cytometry; RNA-seq, RNA-sequencing; sc, single cell; SES-CD, simple endoscopic score for CD.

### Enzymatic digestion

Biopsies were collected in HBSS media (Gibco) containing 2% FCS and 0.2% amphotericin B. The intestinal tissue was transferred to HBSS supplemented with 1 mM DTT (Sigma) and placed on a rolling device for 10 min at 4°C. After discarding the supernatant, the intestinal tissue was transferred to HBSS supplemented with 2% FCS and 5 mM EDTA and shaken (2x) at 180 rpm for 30 min at 37°C. The tissue suspension was passed through a 70µm cell strainer (Costar) and constituted the intraepithelial population. To obtain lamina propria T cells, the remaining intestinal biopsies were subsequently incubated for 1 hour at 37°C with 1mg/ml Collagenase IV (Sigma) in RPMI medium (supplemented with 10% FCS, 100 U/ml penicillin-streptomycin and 0.2% Fungizone), then forcefully resuspended through a 19G needle, washed and filtered with 70 µm cell strainer (Costar). The cell suspensions were used for bulk and single-cell RNA-sequencing after sorting different T cell subsets.

### Fluorescence-activated Cell Sorting

In preparation of fluorescence-activated cell sorting (FACS) the intestinal cells were incubated with the surface antibodies for 20 minutes in supplemented RPMI (2% FCS, 1% penicillin and streptomycin, 0.2% Fungizone) at 4°C, and subsequently washed in FACS buffer before sorting on a FACSAria(tm) III (BD) (for gating strategy see supplemental figure 1A). Antibodies used: fixable viability dye eF506 (65-2860-40), anti-human CD3 eF450 (clone OKT3, 48-0037-42; eBioscience), TCRγδ BV510 (clone B1, 331220), CD3 AF700 (clone UCHT1, 300424), CD4 BV785 (clone OKT4, 317442; Biolegend), CD8α APC-Cy7 (clone SK1, 557834), CD127 BV421 (clone HIL-7R-M21, 562436), CD25 PE-Cy7 (clone M-A251, 557741), CD69 PE (clone FN50, 555531; BD Biosciences), CD8α PerCP-Cy5.5 (clone RPA-T8, 301032), CD127 AF647 (clone HCD127, 351318; Biolegend), TCRγδ FITC (clone IMMU510, IM1571U, Beckman Coulter), CD45RA APC-Cy7 (clone HI100, 2120640), CCR7 APC (clone G043H7, 2366070, Sony Biotechnology). Flow data was analyzed using FlowJo v10.

### Bulk RNA-sequencing

The sorted cells were thawed for TRIzol (Thermo Fisher Scientific) RNA extraction and stored at −80°C until library preparation. Sequencing libraries were prepared using the Cel-Seq2 Sample Preparation Protocol and sequenced as 75bp paired-end on a NextSeq 500 (Utrecht Sequencing Facility). The reads were demultiplexed and aligned to the human cDNA reference genome (hg38) using BWA (version 0.7.13). Multiple reads mapping to the same gene with the same unique molecular identifier (UMI, 6bp long) were counted as a single read.

Raw counts of splice variants were summed and the raw counts were subsequently transformed employing variance stabilizing transformation. Differential analysis was performed using DESeq2 (Wald’s test) with a p-adjusted value < 0.1 considered statistically significant. For visualization purposes the R package DESeq2 was employed. Pathway analysis was performed on the differentially expressed genes as input in Toppfun with standard settings. Gene set enrichment analysis (GSEA^17^ v4.0.3), with as input the normalized data (output DESeq2), was used to assess enrichment of gene sets derived from Magnuson *et al*.^18^, Guo *et al*.^19^ and Mijnheer *et al*.^20^. One thousand random permutations of the gene sets were used to establish a null distribution of enrichment score against which a normalized enrichment score and FDR-corrected *q* values were calculated. Identification of key-regulators was performed using RegEnrich^21^ v1.0.1 based on the differential gene expression data followed by unsupervised gene regulatory network inference (GENIE3) to construct a network based on the raw gene count data, and GSEA was used for enrichment analysis. Network visualization was performed with Cytoscape^22^ v3.9.0.

### Single-cell RNA-sequencing

Live CD3^+^CD8^-^CD4^+^ cells were sorted (for gating strategy see supplemental figure 1A) into 384-well hard shell plates (Biorad) with 5 μl of vapor-lock (QIAGEN) containing 100-200 nl of RT primers, dNTPs and synthetic mRNA Spike-Ins and immediately spun down and frozen to −80°C. Cells were prepared for SORT-seq as previously described.^23^ Illumina sequencing libraries were then prepared with the TruSeq small RNA primers (Illumina) and sequenced single-end at 75 bp read length with 75.000 reads per cell on a NextSeq500 platform (Illumina). Sequencing reads were mapped against the reference human genome (GRCh38) with BWA.

Quality control was performed in R with seurat and cells were dropped when the number of genes was < 150, and/or the percentage of mitochondrial genes was > 35%. Potential doublets were eliminated based on the gene/UMI ratio. Cut-offs were set based on visual inspection of the distribution and preliminary clustering analyses. The raw data expression matrices were subsequently analyzed using Seurat (v4^24–26^) following the outline provided by the distributor (https://satijalab.org/seurat/). Each dataset was log normalized, variable features were determined using vst, and the percent of mitochondrial genes and difference between G1 and G2M phase of the cell-cycle were regressed out. Thereupon, the datasets were merged with Seurat and batch-effect correction for patient was performed with Harmony.^27^

For dimensionality reduction first the PC’s were calculated (RunPCA) and clustering was performed with UMAP (RunUMAP: 30 dimensions, n.neighbors 20; FindNeighbors: clustering resolution of 0.8). Index sort data was analyzed in FlowJo v10 using the Index sort v3 script (https://www.flowjo.com/exchange/#/plugin/profile?id=20). Median Fluorescent Intensity cut-offs were determined based on the gating, and imported as metadata. Subsequent differential gene expression was performed using the MAST test (standard settings, regression for patient) with a p-adjusted value < 0.05 considered statistically significant. Visualization were performed with seurat.

### Imaging mass cytometry

Intestinal biopsies were fixed in 10% neutral buffered formalin, paraffin-embedded, and two slides containing consecutive 4 µm-thick sections of all samples were prepared. One slide was stained with haematoxylin and eosin (HE) for histological assessment and the second slide was stained for imaging mass cytometry (IMC). IMC combines immunohistochemistry with high-resolution laser ablation of stained tissue sections followed by cytometry by time of flight (CyTOF) mass cytometry enabling imaging of multiple proteins at subcellular resolution.^28^ All antibodies were conjugated to lanthanide metals (Fluidigm, San Fransisco, CA, USA) using the MaxPar antibody labeling kit and protocol (Fluidigm), and eluted in antibody stabilization buffer (Candor Bioscience) for storage.

The slide was baked for 1.5 hours at 60°C, deparaffinized with fresh xylene for 20 minutes and subsequently rehydrated in descending grades of ethanol (100% (10 minutes), 95%, 80%, 70% (5 minutes each). After washing for 5 minutes in milliQ and 10 minutes in PBST (PBS containing 0.1% Tween-20), heat-induced epitope retrieval was conducted in Tris/EDTA (10 mM/1 mM, pH 9.5) for 30 minutes in a 95°C water bath. The slide was allowed to cool to 70°C before washing in PBST for 10 minutes. To decrease non-specific antibody binding, tissue sections were blocked with 3% BSA and Human TruStain FcX (1:100, BioLegend) in PBST for 1 hour at room temperature. The antibody cocktail was prepared by mixing all antibodies at concentrations specific for the assay in PBST+0.5% BSA. After careful removal of the blocking buffer, the slide was incubated overnight at 4°C with the antibody cocktail. Antibodies used: E-cadherin 142Nd (metal tag) (clone 24E10, CST3195BF, Cell Signaling Technology), CD3 170Er (polyclonal, A045229-2, Dako), CD4 156Gd (clone EPR6855, ab181724, Abcam), CD8α 162Dy (clone C8/144B, 14-0085-82, Thermo Fisher Scientific), FOXP3 155Gd (clone 236A/E7, ab96048, Abcam), Ki-67 168Er (clone B56, 556003, BD Biosciences), Granzyme B 169Tm (clone D6E9W, CST46890BF, Cell Signaling Technology). Following three 5 minutes washes in PBST and rinsing in milliQ the tissue was counterstained with 0.1% toluidine blue for 5 minutes to enable tissue structure visualization under bright field microscopy if desired. Upon washing for 5 minutes in milliQ, the slide was incubated with Ir-intercalator (1:500 in PBST, Fluidigm) for 60 min at room temperature. Finally, the slide was washed in milliQ and air dried for at least for 20 min at room temperature.

Images were acquired at a resolution of 1 µm using a Hyperion Imaging System (Fluidigm). Regions of interest were selected based on the HE slides after which areas with an approximate size of 1,000 × 1,000 µm were ablated and acquired at 200 Hz. Pseudo-colored intensity maps were generated of each mass channel. Composite images were created and analyzed for each sample using Image J (version 1.47), and any changes to the brightness or contrast of a given marker were consistent across all samples.

### Statistical analysis

Flow cytometric data was analyzed with a two-tailed Mann-Whitney U or Wilcoxon test with a p-adjusted value < 0.05 considered statistically significant. Data were analyzed with GraphPad Prism (GraphPad Software version 7.0, La Jolla, CA, USA). Singe cell and bulk RNA-sequencing was statistically analyzed as described in the relevant paragraphs.

### Data availability statement

The raw count data generated with the bulk and single cell RNA-sequencing are available on https://github.com/lutterl/CD4-T-cells-ileum.

### Ethics statements

The study protocols (TCBio 17/443, 17/444, 18/522 and NL28761.091.09) were approved by the research ethics committee of the University Medical Center Utrecht (Utrecht, The Netherlands) and the Radboud University Nijmegen Medical Centre (CMO Regio Arnhem-Nijmegen, Nijmegen, The Netherlands), respectively. Written informed consent was obtained from each participating patient before any study-related procedure was performed. The procedures were performed in accordance with the Declaration of Helsinki.

## Results

### Increased epithelial and lamina propria regulatory and effector CD4 T cells in (active) ileal Crohn’s disease

We first analyzed the composition of major T cell populations in the epithelium and lamina propria of the human ileum. Therefore, we obtained ileal biopsies from control subjects as well as paired non-inflamed and inflamed ileal biopsies from patients with CD (gating strategy: supplemental figure 1A). We assessed the presence of the following T cell subsets: T cell receptor (TCR)γδ, TCRγδ^-^CD8^-^CD4^-^, CD8, CD4CD8α (of which the vast majority is expected to be CD4CD8αα), effector (CD4^+^CD127^+^CD25^-^), and Treg (CD4^+^CD127^low^CD25^high^) cells (supplemental figure 2A). The non-inflamed of control subjects and patients with CD was found to display a comparable T cell subset composition, except for an increase in Tregs in the lamina propria of patients with CD (*p* = 0.0295). Inflammation resulted in a further increase in Tregs (*p* = 0.0078) and a decrease in CD4CD8αα T cells (*p* = 0.0391) in the lamina propria compared to the paired non-inflamed ileum of patients with CD. The epithelium of the inflamed ileum displayed increased percentages of Tregs and CD4 effector T cells as well (*p* = 0.0156 and 0.0078, respectively), with a concomitant decrease in CD8 T cells (*p* =0.0156). Furthermore, within the CD4 effector T cell subset, an increase in CD69^-^ cells was observed in CD patients (*p* = 0.0451 for lamina propria CD45RA^-^CD69^-^ in CD versus control subjects), which was most pronounced in the inflamed epithelium (*p* = 0.0078 for CD45RA^-^CD69^-^ and CD45RA^+^CD69^-^ in inflamed versus non-inflamed epithelium and lamina propria, supplemental figure 2B), indicating infiltration/expansion of CD69^low^ effector memory and naive CD4 T cells. Thus, most inflammation-associated changes in the overall T cell composition impact the regulatory and migrating/infiltrating CD69^low^ CD4 effector subset content, and occur within the epithelium.

To additionally assess potential changes in CD4 T cell distribution throughout the mucosal compartment, we performed IMC on paired inflamed and non-inflamed ileal tissue derived from two control subjects and three patients with CD (figure 1A). In the inflamed lamina propria of CD patients diffuse CD4 T cell infiltrates were present (figure 1A, upper row), and loss of normal mucosal architecture can be observed (figure 1A, upper row outer right). FOXP3^+^ Tregs were predominantly located within CD4-rich regions in the lamina propria, especially in the inflamed ileum (figure 1A, upper row). Within the ileal mucosa of control subjects no Ki-67^+^ or Granzyme B^+^ CD4 T cells were observed (figure 1A, outer left panels). In patients with CD, the non-inflamed lamina propria contained a few Ki-67^+^ CD4 T cells and Ki-67^+^ Tregs (FOXP3^+^), as well as Granzyme B^+^ CD4 T cells (figure 1A, inner left panels). However, in inflamed tissue a higher ratio of CD4 and FOXP3^+^ CD4 T cells expressing Ki-67^+^ or Granzyme B^+^ was found (figure 1A, right panels), and within the inflamed lamina propria some Ki-67 and Granzyme B double-positive CD4 T cells were observed (figure 1A, right panels). Altogether, these IMC data indicate that CD4 (FOXP3^+^) T cells both proliferate locally and gain cytotoxic effector function in CD patients, especially in the inflamed mucosa.

**Figure 1.**
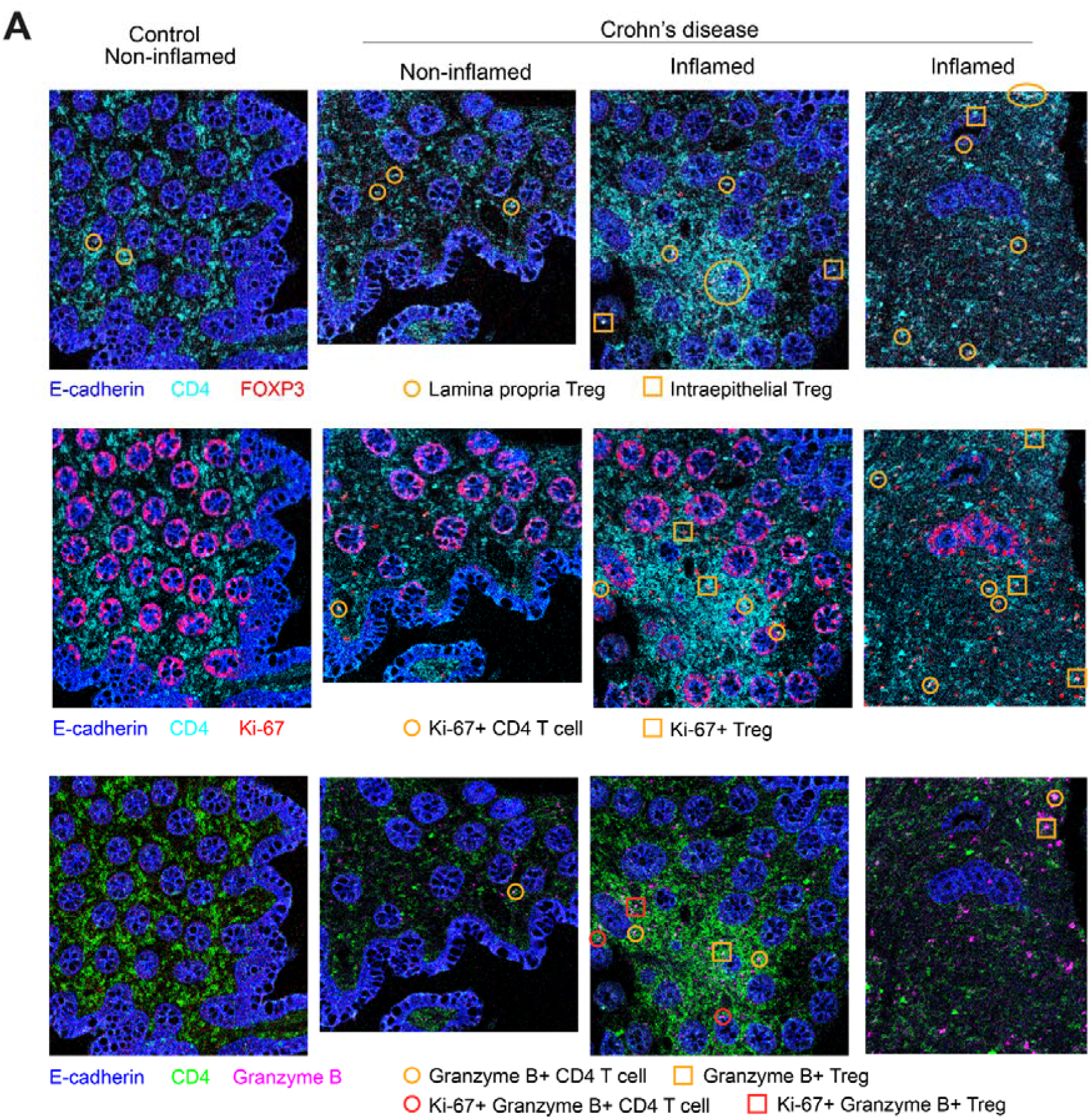
T cell distribution in the human ileum in health and Crohn’s disease. (A) Representative imaging mass cytometry images of the distribution of CD4 and regulatory T cells (upper row), Ki-67 (middle row) and Granzyme B (lower row) in control subjects (outer left), non-inflamed (inner left) and inflamed (right) ileum of CD patients. Control subjects *n* = 2, CD *n* = 3. A few examples are highlighted per image. Each column shows the same image, but colored for different targets per row.

### Inflammation is associated with effector Treg differentiation in the epithelium

We then performed bulk RNA-sequencing of sorted CD4CD8αα T cells, Tregs (CD4^+^CD127^low^CD25^+^), and effector T cells (CD4^+^CD127^+^CD25^-^) subdivided in CD69^-^ and CD69^high^ (Trm cells) from the ileum of control subjects and patients with CD (paired inflamed and non-inflamed) (gating strategy: supplemental figure 1A). Although principal component analysis revealed partial clustering of surface marker-based CD4 T cell subsets, there was no complete distinction of all subsets (figure 2A, left; supplemental figure 3A). For example, lamina propria CD4CD8αα T cells only differed from CD69^high^ Trm cells by expression of *CD8A*. In the epithelium, however, 13 upregulated and 138 downregulated genes were observed in CD4CD8αα T cells compared to CD69^high^ CD4 Trm cells, although none of these genes have clear effector, regulatory nor cytotoxic, functions (supplemental table 1). In mouse epithelium, T-bet and Runx3 induce the intraepithelial CD4 T cell program including CD8α expression.^29^ Both T-bet and Runx3 were, however, not differentially expressed in human small intestinal CD4CD8αα T cells in the epithelium compared to the lamina propria, nor compared to CD69^high^ Trm cells or Tregs. CD4CD8αα T cells did differ from Tregs with downregulated *FOXP3*, and downregulation of the canonical Treg markers *IL2RA, IKZF2, TIGIT* and *TNFRSF18* (supplemental table 1). This is in line with previous data^15,30^ and indicates that CD4CD8αα T cells are clearly different from mouse and human small intestinal FOXP3^+^ Tregs. As expected, the Trm signature was more enriched in CD69^high^ Trm compared to CD69^-^ CD4 T cells within the lamina propria (supplemental figure 3B). Among the 395 upregulated and 31 downregulated genes were many known Trm genes (e.g. *CD69, CXCR6, IFNG* and *ITGA1*^4^) (supplemental table 1).

**Figure 2.**
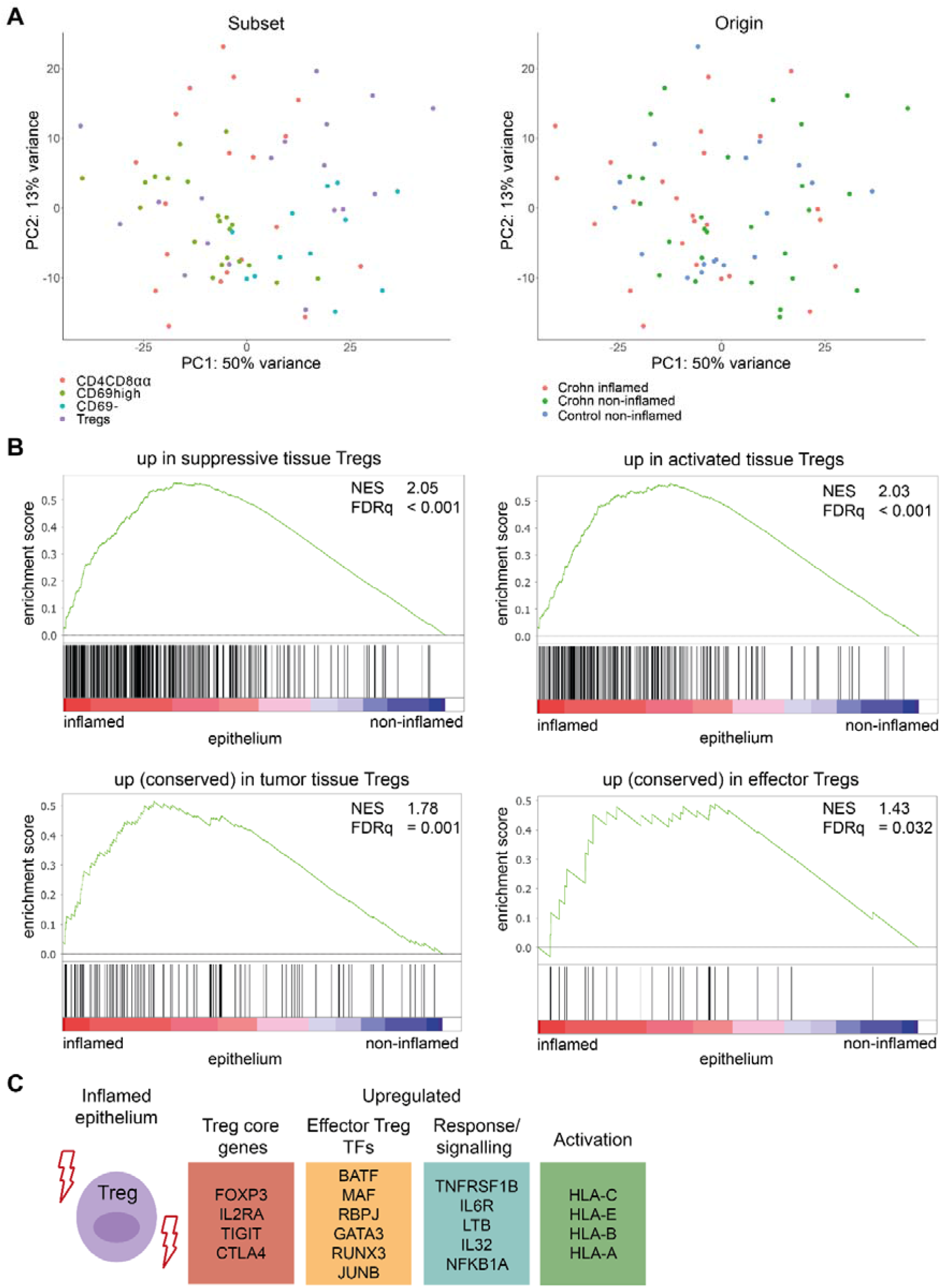
Effector Treg differentiation in the inflamed epithelium. (A) Unsupervised principal component analysis of all CD4 T cell subsets analyzed by bulk RNA-sequencing colored on subset (left; pink = CD4CD8αα T cells, green = CD69^high^ Trm cells, blue = CD69^-^ T cells, purple = Tregs), and status (right; pink = CD inflamed, green = CD non-inflamed, blue = control non-inflamed). (B) Gene set enrichment analysis of suppressive^19^ (upper left), activated^19^ (upper right), tumor-derived tissue Treg^18^ (lower left) and conserved human effector Treg^20^ (lower right) signatures in pairwise comparisons of Tregs derived from non-inflamed and inflamed ileum of CD patients, represented by the normalized enrichment score (NES) and FDR statistical value (FDRq).

Moreover, the origin of the samples (control subjects or patients with CD) was not found to fully drive clustering either, irrespective of the presence of inflammation (figure 2A, right; supplemental figure 3A). First, we analyzed the impact of inflammation on the whole CD4 T cell population (combined CD69^high^ Trm cells, CD4CD8αα T cells and Tregs). Whereas only 1 differentially expressed gene was found when comparing CD4 T cells from paired inflamed and non-inflamed lamina propria of patients with CD, comparing CD4 T cells from inflamed lamina propria with healthy non-inflamed control subjects resulted in 1030 upregulated and 822 downregulated genes (supplemental table 1). Upregulated genes included *STAT1, BATF, PRDM1, GZMB, IL22* and *CCR6*, and associated with processes related to cell activation, cell cycling, and stress. In the epithelium, significant differences in inflamed versus non-inflamed epithelium of patients with CD were found (1009 upregulated and 58 downregulated genes, supplemental table 1). Here, CD4 T cells from the inflamed epithelium showed, among others, increased translation and protein transport, T cell activation and responses to IFNγ, with upregulated genes including ribosomal genes, *CXCR3, CCR7, TOX2* and *KLF2*.

Even though the relative composition within the CD4 T cell population of the inflamed ileum shifted (supplemental figure 2A,B), only a limited number of transcriptional changes (<10) was found for any of the sorted bulk CD4 T cell subsets (CD69^-^, CD69^high^, CD4CD8αα T cells and Tregs) derived from inflamed ileum, except for intraepithelial Tregs. Intraepithelial Tregs from inflamed ileum mucosa acquired an effector Treg profile with upregulation of *FOXP3, TIGIT* and *TNFRSF1B*, among others (figure 2 B,C, supplemental table 1). Enrichment of gene signatures procured from literature for suppressive, activated, tumor suppressive Tregs and a conserved effector Treg program in inflammation-derived intraepithelial Tregs corroborate these observations (figure 2B).^18–20^ *FOXP3, GATA3, BATF, RUNX3* and *MAF* were among the upregulated transcription factors (TFs) in Tregs from the inflamed epithelium which have been associated with effector Treg differentiation.^20,31–33^ Both GATA3 and RUNX3 are crucial in suppression of inflammation by Tregs.^32,33^ Thus, the most pronounced transcriptional change in CD4 T cell subsets in CD inflammation is the strong effector Treg differentiation in the epithelial compartment.

### Mucosal subcompartment drives the CD4 T cell transcriptomic profile in the ileum

Clear separation of CD4 T cell subsets was observed based on compartment of residence; the epithelium and lamina propria of the ileum (figure 3A, supplemental table 1). Gene ontology analysis revealed that CD69^high^ Trm cells in the epithelium were enriched for pathways associated with activation, T cell differentiation and mRNA processing (790 upregulated and 299 downregulated genes for epithelium versus lamina propria) (figure 3B). However, pathways related to activation were upregulated in both lamina propria Tregs and CD4CD8αα T cells (708 and 239 upregulated and 1109 and 734 downregulated genes for epithelium versus lamina propria, respectively). These included cell activation and T cell differentiation for CD4CD8αα T cells, and mRNA processing, stress response and cell cycle regulation for Tregs (figure 3B). The enriched pathways for the different CD4 T cell subsets were predominantly regulated via different genes (21 shared genes in the epithelium and 22 in the lamina propria). In the lamina propria, *TRAF1, TRAF4* and *BIRC3* involved in TNF(R) signaling, as well as *IL4R* and *IL2RA* associated with T cell activation and differentiation, were shared among all CD4 T cell subsets^34^, whereas in the epithelium genes involved in migration and motility were shared (e.g. *CLIC1*^35^, *GNAQ*^36^, *FAM107A*^37^) (supplemental table 1).

**Figure 3.**
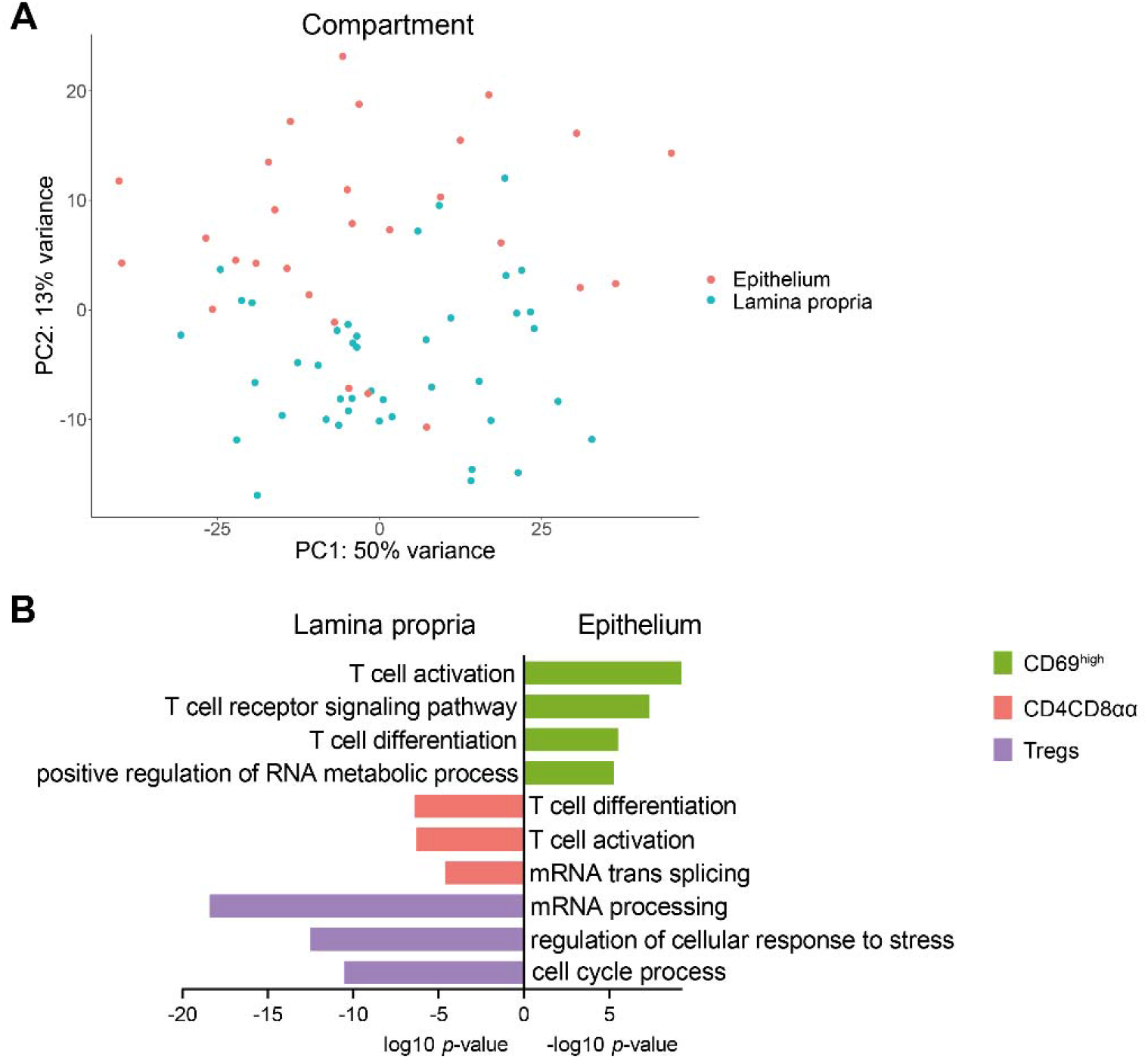
Mucosal subcompartment shapes the transcriptomic profile of CD4 T cells. (A) Unsupervised principal component analysis of all CD4 T cell subsets analyzed by bulk RNA-sequencing colored on compartment of residence (pink = epithelium, blue = lamina propria). (B) Selected biological process terms related to the upregulated genes in the lamina propria for CD69^high^ Trm cells (green), and in the epithelium for CD4CD8αα T cells (pink) and Tregs (purple). (C) Selected genes upregulated in Tregs in the inflamed epithelium. TFs, transcription factors.

Subsequently, we performed single-cell RNA-sequencing of paired non-inflamed and inflamed ileum-derived CD4 T cells (CD3^+^TCRγδ^-^CD4^+^) from the epithelium and lamina propria of patients with CD to assess potential heterogeneity missed with bulk RNA-sequencing. These data supported our observation that compartment is the primary driver of CD4 T cell clustering (figure 4A-C, gating strategy: supplemental figure 1A). Unsupervised clustering of 1573 cells after quality control revealed five CD4 T cell clusters represented in every patient (figure 4A,D). Cluster 1 (18.6% of allcells) was characterized by *KLF2, SELL, CCR7, TCF7* and *LEF1*, and thus resembled recently migrated/recirculating T cells (figure 4A). This cluster contained both intraepithelial and lamina propria T cells (figure 4E). Similarly, cluster 2 (14.3% of all cells) comprised both lamina propria and intraepithelial cells constituting Tregs (*FOXP3, TIGIT, CTLA4, IKZF2* and *BATF*) (figure 4A,E). Cluster 3 and 4 predominantly comprised lamina propria-derived CD4 T cells (figure 4E). Cluster 3 (26.8% of all cells) seemed to contain a mix of Th1- and Th17-skewed cells identified by *CCL20, NFKB1/2, TRAF1* and *BHLHE40*, whereas cluster 4 (11.1% of all cells)showed higher expression of heat shock protein family members suggesting that these cells were stressed (figure 4A). The last cluster (cluster 5, 29.2% of all cells) was comprised of intraepithelial CD4 T cells distinguished by expression of innate and/or cytotoxic related genes including *FOS(B), EGR1, GZMA, JUN* and *IER2* (figure 4A,E, supplemental table 2). CD4CD8αα T cells mixed with the (resident) CD4 T cells and not with the Tregs, in line with the bulk RNA-sequencing data (figure 4C). Also, in accordance with the bulk data, inflammation did not result in distinct CD4 T cell profiles, although, congruent with the protein data, we observed a relative increase in CD69^-/low^ CD4 T cells in all defined clusters in inflammation (figure 1B, 3F). Almost 50% of cluster 1 and 2 cells comprised CD69^low^ T cells, compared to approximately 10-15% for the other clusters, and these clusters showed the highest relative increase of CD69^low^ T cells in inflamed tissue. In summary, bulk and single-cell RNA-sequencing data revealed that the intra-tissue compartmentalization of CD4 T cells is the primary driver of their transcriptomic landscape. During inflammation, the expression profiles of CD4 Trm cells is largely preserved, with a significant influx of CD69^low^ migrating CD4 T cells.

**Figure 4.**
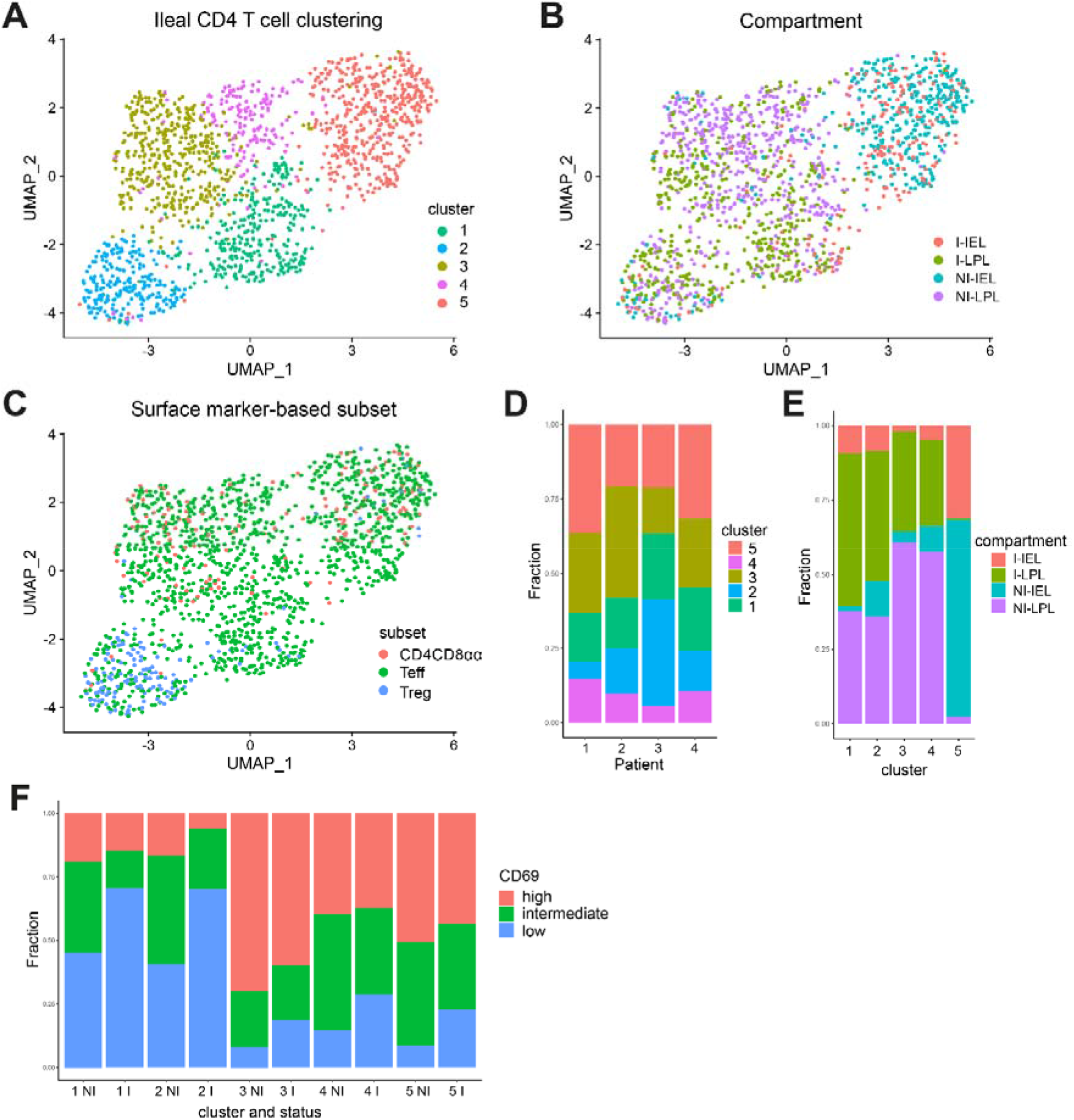
CD4 T cell clusters in the paired non-inflamed and inflamed ileum. (A) Dimensionality reduction of all CD4 T cells (CD3^+^TCRγδ^-^CD4^+^) from the epithelium and lamina propria of four patients with Crohn’s disease. Cells are colored based on the assigned cluster. (B) As per A colored on the compartment (IEL = intraepithelial T cell, LPL = lamina propria T cell) and status (I = inflamed, NI = non-inflamed). (C) As per A colored on surface marker-based identification. CD4CD8αα T cell = CD4^+^CD8α^+^ (pink), effector T cell (Teff) = CD4^+^CD127^+^CD25^-^ (green), Treg = CD127^-^CD25^+^ (blue). (D) Reproducible composition of the CD4 T cells across the four included patients. y-axis: fraction of cells colored on the cluster as shown in A and separated per patient on the x-axis. (E) Composition of the compartment (epithelium/lamina propria) and status (non-inflamed/inflamed) origins per cluster. y-axis: fraction of cells colored on compartment and separated per cluster as shown in A on the x-axis. (F) Composition of CD69 protein expression among non-inflamed (NI) and inflamed (I) derived CD4 T cells for all clusters, separated by low (negative, blue), intermediate (green) and high (pink) CD69 expression. y-axis: fraction of cells colored on CD69 expression, and separated per cluster, and non-inflamed/inflamed ileum.

### The chromatin remodeling genes ARID4B and SATB1 are predicted key regulators of lamina propria to epithelium translocation of CD4 T cells

Since we observed differences in the transcriptional profile of CD4 T cells derived from the lamina propria and epithelium we performed a data-driven network and enrichment analysis (RegEnrich^21^). Herewith, we could identify which key gene regulators, based on TFs and co-factors, were involved in lamina propria to epithelial translocation and adaptation (combined CD69^high^ Trm cells, CD4CD8αα T cells and Tregs), irrespective of inflammation. Top predicted regulators enabling translocation of CD4 T cells to the epithelial compartment were *ARID4B, SATB1, CREM, BAZ1A* and *NFKB1* (all negative, i.e. regulators upregulated in the lamina propria and downregulated in the epithelium) (figure 5A, supplemental table 3). Most regulators were found to be downregulated. Of interest, downregulation of many of the abovementioned regulators are involved in chromatin remodeling and inhibit T cell functioning. The negative regulator ARID4B is known as a master regulator of the phosphatidylinositol-3-kinase (PI3K) pathway related to T cell activation and proliferation.^38,39^ Downstream target genes of the downregulated SATB1 are also involved in T cell effector functions including *TRAF1* and *TRAF4* (TNF receptor-associated factors), *NFKB*, interleukin receptors (e.g. *IL4R, IL12RB2*) and *IL2*. Additionally, among the positive regulators were *ELF4, TRIM21* and *TRIM27*, all linked to negative regulation of T cell function. *ELF4* has been associated with CD8 T cell quiescence and suppressed proliferation and downregulation of CCR7, CD62L and KLF2 and thereby tissue retention.^40^ *TRIM21* (Ro52) has been linked to negative regulation of pro-inflammatory cytokine production^41^ by limiting Th1/Th17 differentiation and thereby suppressing tissue inflammation in an IL23/Th17-dependent manner.^41,42^ Lastly, *TRIM27* does not inhibit Th-cell differentiation but negatively impacts CD4 T cell functioning via the PI3K pathway.^43^ Furthermore, positively regulator-associated genes encompassed *FASLG* and programmed cell death genes, whereas negatively associated downstream genes included cell cycling genes. This suggests a different regulation of proliferation and cell death in these compartments. Altogether, the predicted network of TF regulators involved in translocation/adaptation of CD4 T cells from the lamina propria to the epithelium showed downregulation of chromatin remodeling, dampening of T cell activation and effector function, as well as regulation of programmed/activation-induced cell death and arrest of cell cycling independent of inflammation. Overall, these data suggest that upon translocation of CD4 T cells to the epithelium their function and lifespan are tightly regulated.

**Figure 5.**
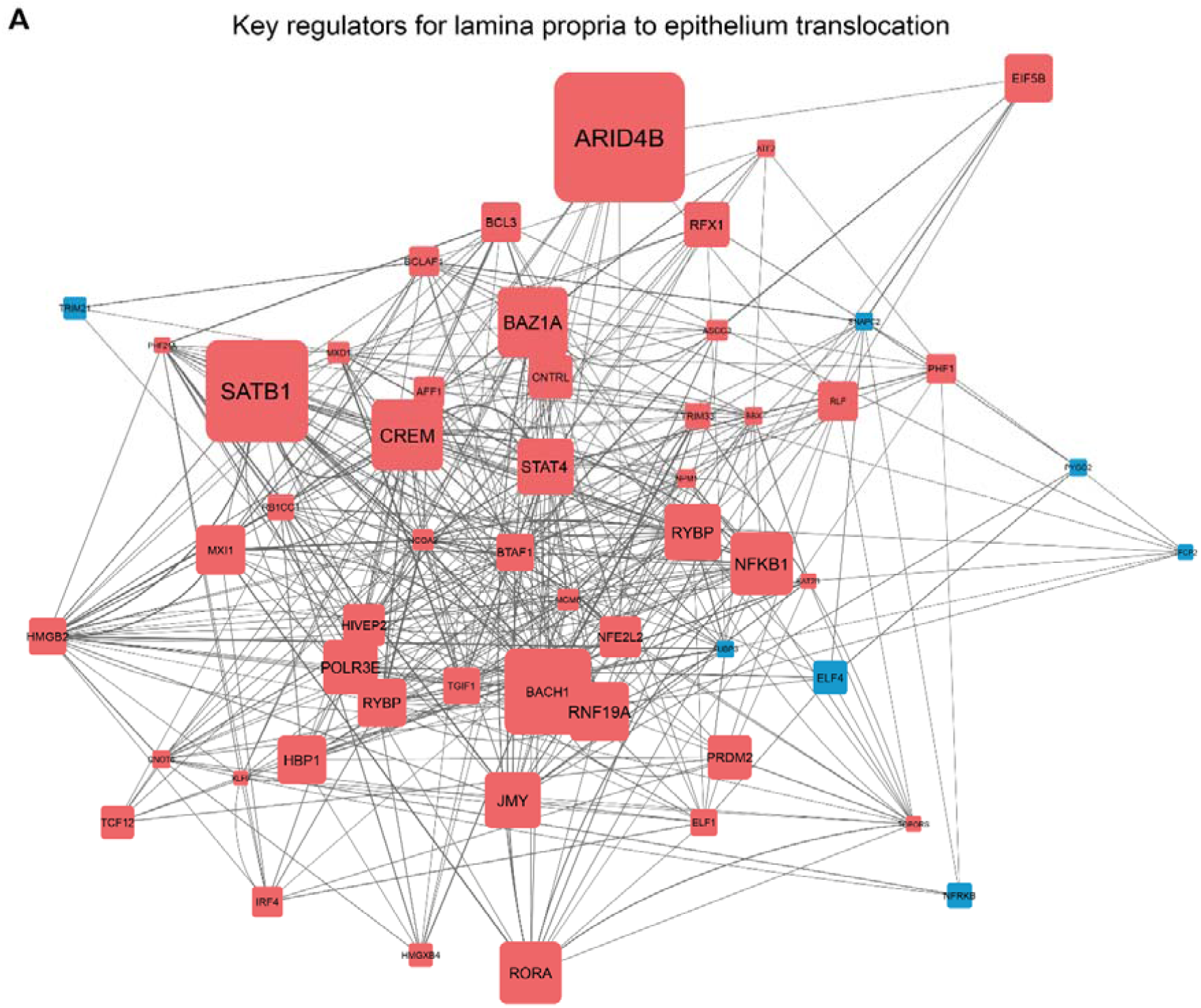
Predicted key regulators of lamina propria to epithelium translocation for CD4 T cells. (A) Network inference of key-regulators driving lamina propria to epithelium CD4 T cell translocation on RNA level, based on unsupervised gene regulatory network analysis followed by gene set enrichment analysis of the transcription factors (TFs) and co-factors (pink = upregulation, blue = downregulation). The grey lines indicate connections between regulators (TFs) and their downstream targets (only TFs are shown), the line thickness represents the correlation weight (thicker = higher correlation). Square size indicates −log10(*p*) for each comparison, with the *p*(-value) derived from differential expression analysis (log2 fold change > 0.5 and *p*-adjusted value < 0.1), text size represents the RegEnrich score; for both, larger indicates higher scores (supplemental table 3).

## Discussion

In the present study, we demonstrate that the transcriptional profile of CD4 effector, Trm cells, CD4CD8αα T cells and Tregs is primarily determined by the compartment of residence (epithelium or lamina propria), irrespective of inflammation. The relative composition of the mucosal CD4 T cell population changes in patients with CD with an increase in Tregs. Upon inflammation, Tregs as well as CD4 naive and migrating/infiltrating CD69^low^ T cells further increase, while the number of CD4 Trm cells, CD8 and CD4CD8αα T cells relatively decrease. These changes are most pronounced in the epithelium. Inflammation induces only few transcriptomic changes in lamina propria CD4 Trm cells, Tregs and CD4CD8αα T cells on the bulk and single-cell RNA-sequencing level, indicating that reshaping of CD4 T cell subset transcriptomes in the inflamed lamina propria of patients with CD is limited. In the inflamed epithelium, however, a considerable upregulation of cell activation, cell cycling and stress response is observed, with Tregs gaining an effector Treg profile, which is also seen in tissues in other inflammatory diseases.^20^ Translocation of CD4 T cells from the lamina propria to the epithelium, irrespective of inflammation, is driven by chromatin remodeling via *ARID4B* and *SATB1*and the main consequence of translocation of these cells to the epithelium, seems to be a dampening of broad (pro-inflammatory) T cell responses.

Previous studies have shown that T cell subsets derived from the small intestine and colon are transcriptionally different, independent of inflammation.^44^ Additionally, research has indicated that there are less differences between T cell subsets (CD4 and CD8 effector T cells, and Tregs and CD4 conventional T cells)^45,46^ and inflamed and non-inflamed gut^47^ than there is between the lamina propria and epithelium.^45–47^ Our study is the first that explores the effects of both the compartment (epithelium or lamina propria) and presence of inflammation in a spectrum of different CD4 T cell subsets and concludes that compartment is the primary driver of the transcriptomic profile of a human small intestinal CD4 T cell.

Our data further indicates that the an increase in migrating/infiltrating CD69^low^ CD4 T cells constitutes a major change in the local T cell population during inflammation in CD. In recent years, the therapeutic armamentarium of IBD has been extended to the anti-integrin therapies vedolizumab (anti-integrin α4β7) and etrolizumab (anti-integrin β7). Sphingosine 1 phosphate (S1P), CCR9 and MAdCAM-1 targeted therapies are also under investigation in clinical trials for IBD.^48^ Based on our data, it might be beneficial to implement lymphocyte trafficking interfering strategies early in the disease to prevent migration/infiltration of these cells. This might also prevent (pathogenic) infiltrating cells to become Trm cells over time.

The transcriptomic differences of CD4 T cells between the epithelium and lamina propria could be due to a preprogrammed destination of T cells or to local factors inducing a compartment-specific profile in the same cell of origin. A recent study in mice showed that T cells receive compartment-specific imprinting for the lamina propria or epithelial compartment in the mesenteric lymph nodes.^46^ However, another study in mice showed that transfer of lamina propria-derived T cells will lead to relocation of these cells to both the lamina propria and epithelium.^49^ Furthermore, overlap in the TCR repertoire for lamina propria and epithelium CD8 Trm cells has been observed,^14^ and several studies have shown that T cells adapt to the local microenvironment.^31,50^ It is likely that both lymph node originated compartment-specific imprinting and the local environment contribute to the transcriptomic profile of T cells. Nevertheless, the extensive transcriptomic differences due to compartment of residence emphasizes the importance of considering the lamina propria and epithelium as separate compartments when deciphering the pathogenic mechanisms of chronic mucosal inflammation. This distinct compartmentalization of CD4 T cells also suggests that maintaining and restoring mucosal integrity is of key importance to prevent inappropriate immune responses.

The predicted key regulators of CD4 T cell lamina propria-to-epithelium translocation included TRIM27 involved in limiting Th1/Th17 differentiation.^41,42^ The set of predicted regulators, among others, could explain the less pronounced Th-cell and more pronounced innate/cytotoxic profile of intraepithelial compared to lamina propria CD4 T cells. Furthermore, both the IMC and bulk RNA-sequencing data indicate that cell cycling/death in the epithelium is tightly controlled, and that local CD4 T cell expansion during inflammatory responses is primarily observed in the lamina propria. Uncontrolled cell death and extensive local expansion in the epithelial layer could potentially disrupt the single-cell layer of epithelial cells and consequently impact mucosal barrier integrity. Altogether, these data indicate a tight regulation of broad (and untargeted) effector function and cell cycling of intraepithelial CD4 T cells, potentially to prevent destructive tissue responses while allowing targeted action via cytotoxicity. That many predicted key regulators involved in the translocation of CD4 T cells from the lamina propria to the epithelium are chromatin remodelers suggests that the adaptation to the microenvironment is imprinted in these cells. Future (single-cell) epigenomic/chromatin accessibility sequencing (e.g. ATAC-sequencing) could help to elucidate the specific chromatin alterations that occur, and thus the compartment-specific imprinting that ensues.

Active disease in patients with CD does not result in transcriptional changes in CD4 T cells within the lamina propria. However, compared to the non-inflamed lamina propria of control subjects, cell activation and stress responses were observed. This strongly suggests that in established CD there are changes in the transcriptome of CD4 T cells across the mucosa. Future research should include newly diagnosed patients or prediagnostic samples (with the latter being hard to collect) to assess how CD4 T cells at baseline or before diagnosis are altered. This could be pre-onset changes as shown by presence of an IBD-like microbiome in healthy co-twins at risk of developing IBD,^51^ the influence of (previous) administered medication as shown in the blood of IBD patients,^52^ or the effect of previous flares. The fact that CD patients with an ileocecal resection often experience relapses at the anastomosis^53^ suggests that the gut mucosa of CD patients exhibits (pre-existent) changes, although the occurrence of relapses could also be (partly) caused by non-immune cell related factors.

Our data indicate that the inflamed epithelium undergoes extensive changes with an upregulation of protein translation and activation of CD4 T cells in patients with CD. The limited transcriptional differences of ileal CD4 Trm cells upon inflammation suggest that infiltrating CD4 T cells might have a pathogenic role. Of note, absence of subset-specific transcriptional changes for CD4 (Trm) cells does not exclude changes on the protein level. For example, our IMC data revealed lamina propria Tregs present in CD4 T cell infiltrates to gain Granzyme B expression and thus cytotoxic effector function which could aid in control of inflammatory responses. Intraepithelial Tregs, however, did adapt by gaining an effector Treg profile. This indicates that the local environment at the epithelial border changes significantly. Tregs are known to suppress pro-inflammatory and/or regulate anti-inflammatory functioning of CD8 T cells, TCRγδ T cells and innate lymphoid cells,^54,55^ which reside in high abundance at the epithelial border.^2,3^ Additionally, Tregs have direct tissue-specific functions including aiding tissue repair^56^ and/or via interaction with epithelial stem cells^57^ thereby preserving/stimulating mucosal barrier integrity. The significant changes observed in the inflamed epithelium suggest that promoting (effector) Treg differentiation and expansion in the gut mucosa as therapeutic strategy might be worthwhile. Indeed, a phase 1/2a clinical trial has shown that injection of antigen-specific Tregs in patients with CD resulted in a significant reduction of disease activity.^58^

CD4CD8αα T cells are commonly referred to as cytolytic T cells derived from CD4 T cells that lost ThPOK and gained RUNX3 and T-bet expression, and are primarily found in the (small intestinal) epithelium of mice.^29,49^ Murine small intestinal intraepithelial CD4CD8αα T cells aid in controlling local inflammatory responses via IL-10 and TGFβ.^15,30^ Human intestinal CD4CD8αα T cell-studies are limited, with one study showing a potential regulatory role via IL-10, CTLA4, GITR, CD25 and LAG3 in the colonic lamina propria.^16^ Our data revealed no difference in the relative presence of ileal CD4CD8αα T cells between the epithelium and lamina propria, or between inflamed and non-inflamed mucosa. Furthermore, downregulation of *ZBTB7B* (encoding ThPOK) was not observed in CD4CD8αα T cells. Our study suggests that CD4CD8αα T cells in the human ileum are (very similar to) Trm cells, and do not express a clear cytotoxic or regulatory transcriptional profile.

In conclusion, our data reveals that there is a CD4 T cell compartment-specific imprinting, with shared key regulators driving translocation from the lamina propria to the epithelium. Epithelial translocation seems to result in an overall dampening of effector function and tight regulation of the lifespan. Furthermore, inflammation in patients with CD results in a relative increase of migrating/infiltrating CD69^low^ CD4 T cells, CD4 T cell activation and IFNγ responses as well as strong effector Treg differentiation within the epithelium. These data indicate that the lamina propria and epithelium of the human ileum are differentially regulated in both control subjects and CD patients, with a pronounced regulatory role in the inflamed epithelium. Our findings suggest that we should consider both compartments in the therapeutic management of IBD.

## Acknowledgements

The authors thank all participants. They thank Herma Fidder, Fiona van Schaik, Meike Hirdes, Joren ten Hove and Auke Bogte from the department of Gastroenterology and Hepatology, UMC Utrecht for help with collecting patient material, Vincent Meij from the department of Surgery, UMC Utrecht for help with collecting pilot patient material, Michal Mokry, Nico Lansu and Noortje van den Dungen for providing bulk RNA-sequencing services, Single Cell Discoveries for providing single-cell RNA-sequencing services, Domenico Castigliego for help with intestinal tissue slide preparation and Yvonne Vercoulen and Mojtaba Amini for providing imaging mass cytometry services.

