## Supplemental data for "Compartment-driven imprinting of intestinal CD4 (regulatory) T cells in inflammatory bowel disease and homeostasis"

**Supplemental data Lutter et al**

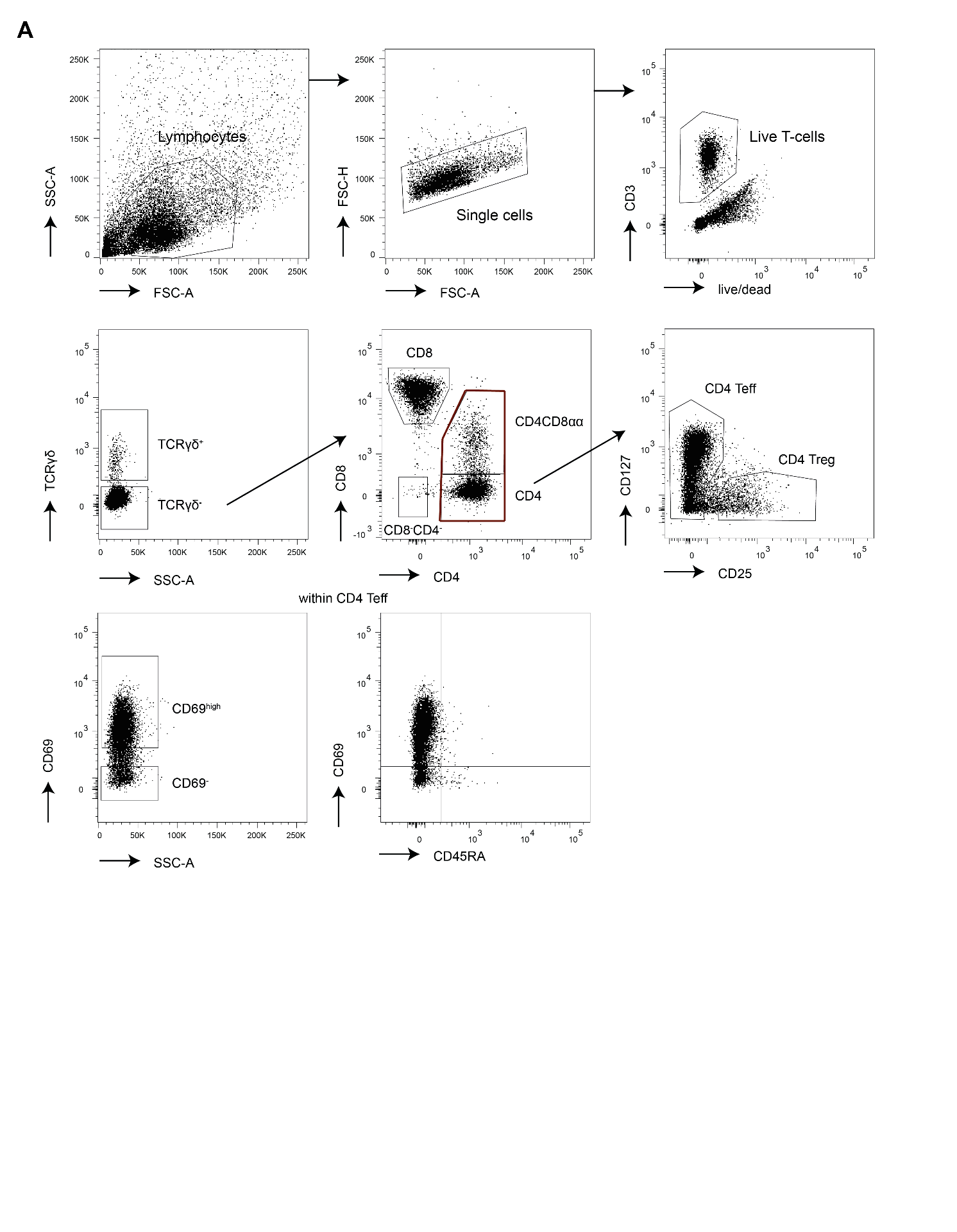

Supplemental figure 1. **Gating strategy.** (A) Gating strategy employed to determine the T cell subset composition, for sorting of CD4^+^CD69^high^, CD4^+^CD69^-^, CD4^+^ Treg, CD4CD8αα (CD8α^+^, vast majority is CD8αα) T cells for bulk RNA-sequencing and CD4^+^ T cells (red outline) for single-cell RNA-sequencing, and to determine the CD69/CD45RA composition. TCR, T cell receptor; Teff, T-effector cells; Treg, regulatory T cells.

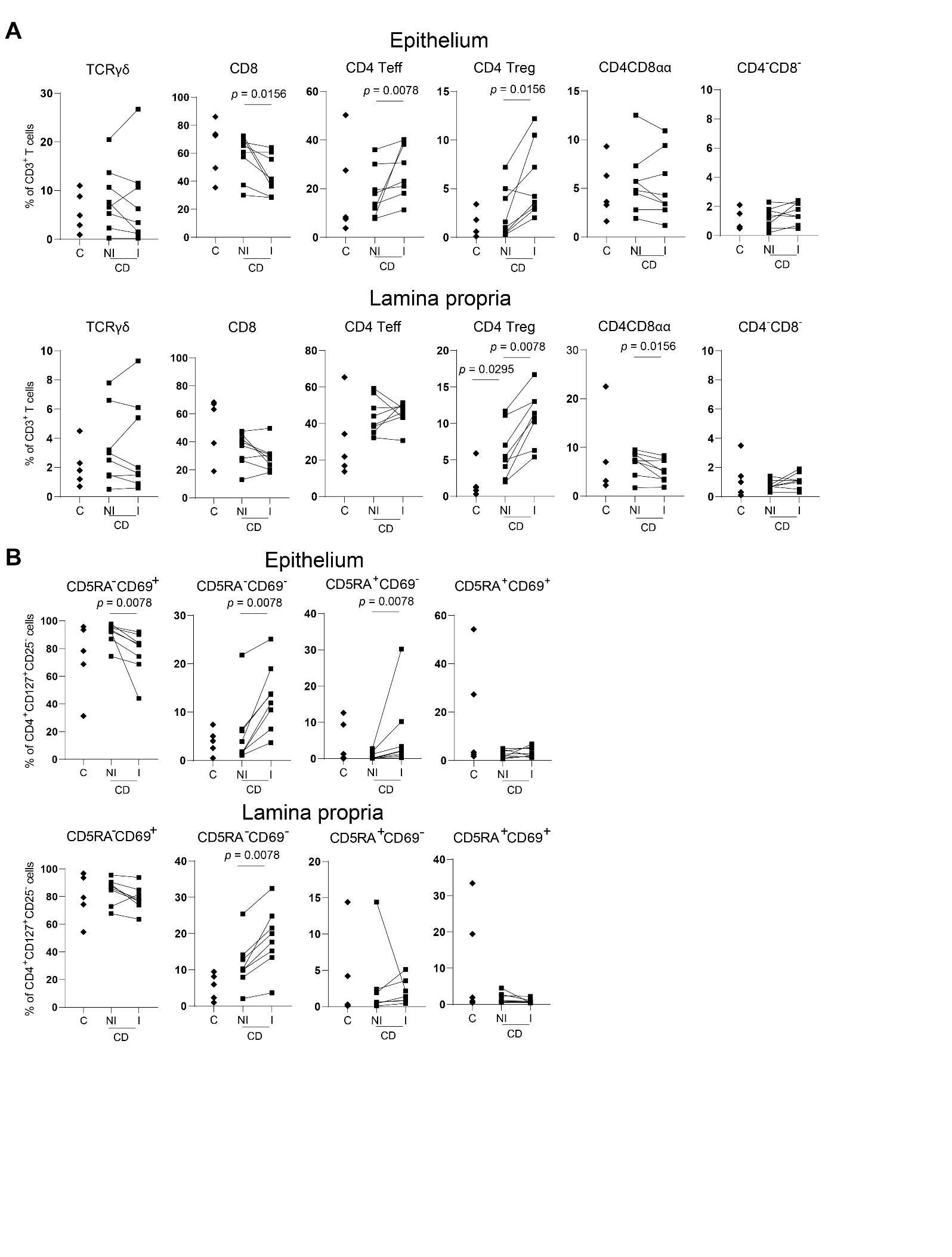

Supplemental figure 2. **T cell composition of the human ileum in health and Crohn’s disease.** (A) Relative composition of T cell receptor (TCR)γδ, CD8, CD4 effector (Teff, CD127^+^CD25^-^), CD4CD8αα (CD4^+^CD8α^+^), CD4 regulatory T cells (Tregs, CD127^low^CD25^high^) and TCRγδ^-^CD8^-^CD4^-^ cells in the epithelium (upper row) and lamina propria (lower row) of control subjects (C), non-inflamed (NI) and inflamed (I) ileum of Crohn’s disease (CD) patients. Lines connect the paired non-inflamed and inflamed datapoints from the patients with CD. Control subjects *n* = 5, CD *n* = 8. (B) Relative composition within CD4 effector T cells of tissue-resident memory (CD45RA^-^CD69^+^), memory (CD45RA^-^CD69^-^), CD4 naive (CD45RA^+^CD69^-^) and recently activated (CD45RA^+^CD69^+^) T cells, similar as to A. Control subjects *n* = 5, CD *n* = 8. Comparisons were performed with a two-tailed Mann Whitney U test for control subject vs non-inflamed CD and Wilcoxon test for paired non-inflamed vs inflamed CD.

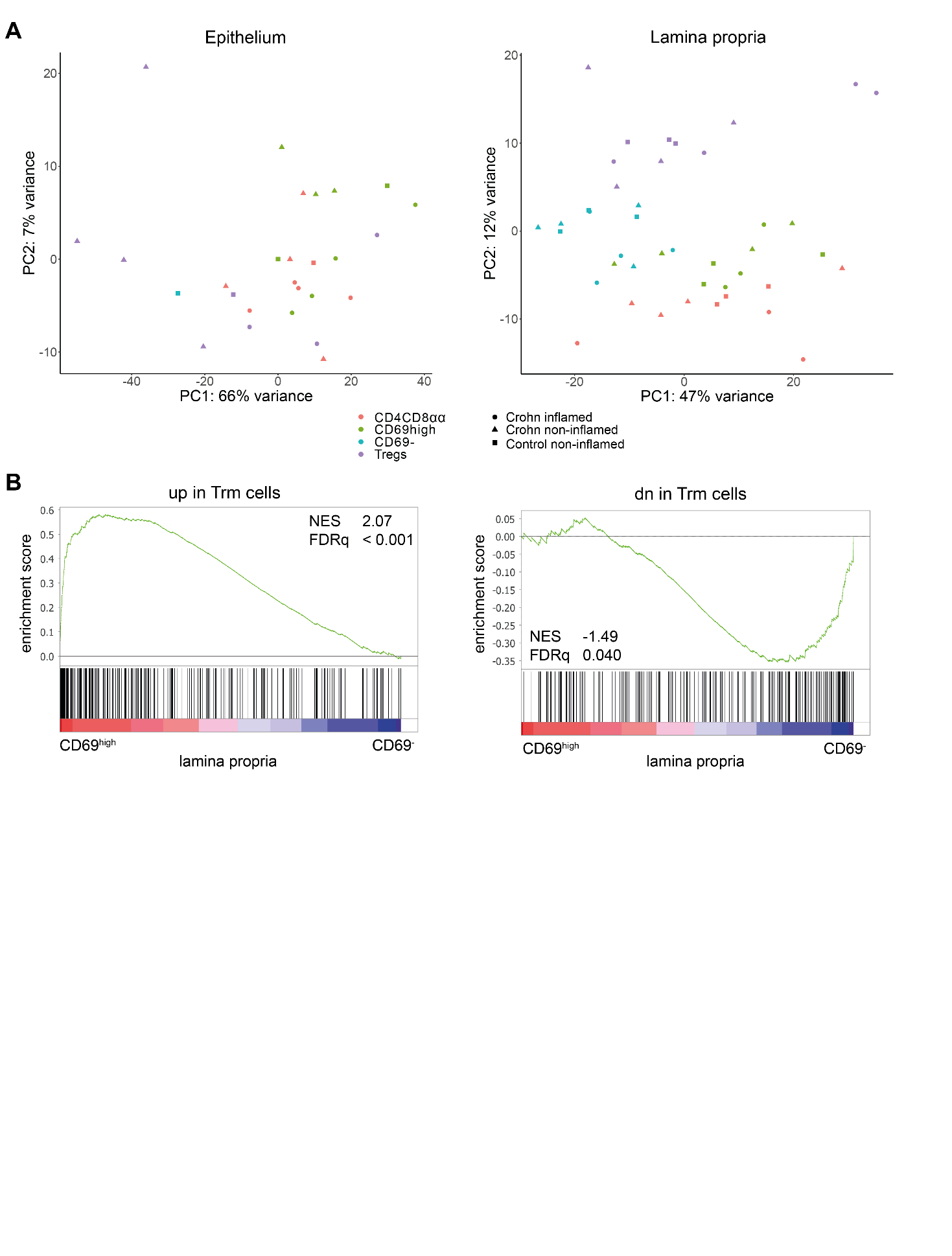

Supplemental figure 3. **Clustering of CD4 T cell subsets in the epithelium and lamina propria.** (A) Unsupervised principal component analysis of all sorted CD4 T cell subsets analyzed by bulk RNA-sequencing, split for epithelium (left) and lamina propria (right), colored on CD4 T cell subset (pink = CD4CD8αα T cells, green = CD4 CD69^high^ Trm cells, blue = CD4 CD69^-^ T cells, purple = Tregs) and status (circle = CD inflamed ileum, triangle = CD non-inflamed ileum, square = control non-inflamed ileum). (B) Gene set enrichment analysis of a Trm signature^4^ (CD69^+^ vs CD69^-^ T cells) with genes upregulated (left) and downregulated (right) in this signature in pairwise comparisons involving transcriptome data of CD69^high^ and CD69^-^ CD4 T cells derived from non-inflamed ileum of control subjects and non-inflamed and inflamed ileum of patients with CD, represented by the normalized enrichment score (NES) and FDR statistical value (FDRq, multiple hypothesis testing using sample permutation).
